# Identification of heterokaryon incompatibility genes in *Aspergillus fumigatus* highlights a narrow footprint of ancient balancing selection

**DOI:** 10.1101/2022.11.25.517501

**Authors:** Ben Auxier, Jianhua Zhang, Francisca Reyes Marquez, Kira Senden, Joost van den Heuvel, Duur K. Aanen, Eveline Snelders, Alfons J.M. Debets

## Abstract

In fungi, a phenomenon termed heterokaryon incompatibility restricts hyphal fusion to occur within an individual since fusion between individuals leads to cell death. Generally, the genes involved are found to be under balancing selection from negative frequency dependent fitness. Here, we assess this in *Aspergillus fumigatus*, a human pathogenic fungus with an extremely high crossover rate. Using auxotrophic markers we screened sexual progeny for compatibility to identify genes involved in this process, the so-called *het* genes. In total, 5/148 (3.4%) offspring were compatible with a parent and 166/2142 (7.7%) sibling pairs were compatible, consistent with several segregating incompatibility loci. Genetic mapping resulted in five loci, four of which could be fine mapped to individual genes, of which we tested three through heterologous expression, confirming their causal relationship. Surprisingly, a population-level analysis of two available independent datasets did not show an increase in Tajima’s D near these loci, normally a hallmark of balancing selection. However, analysis of closely related species did show trans-species polymorphisms across >10 million years, and equal allele frequencies within *A. fumigatus*. Using available *de novo* assemblies, we show that these balanced polymorphisms are restricted to within several hundred base pairs flanking the coding sequence, potentially due to this species’ high crossover rate. In addition to identifying the first *het* genes in an *Aspergillus* species, this work highlights the interaction of long-term balancing selection with a high recombination rate. Future mechanistic work on these *het* genes may provide novel routes for clinical therapies, as well as opportunities for strain improvement in biotechnology.

## Introduction

The ability to differentiate self from nonself is a ubiquitous factor of multicellular life. Perhaps best studied in mammalian MHC loci, such processes prevent fusion between genetically distinct organisms, in effect determining what constitutes an individual (Billingham et al. 1953; Boehm 2006). This ability to distinguish self from nonself is also important for fungi. During the growth of a fungal colony, it benefits from fusions within its own mycelial network (Bastiaans et al. 2015). However, fusion between mycelia of different individuals allows for the propagation of selfish genetic elements such as viruses (Anagnostakis 1983), plasmids and mitochondria (Debets et al. 1994), and nuclei (Bastiaans et al. 2016; Grum-Grzhimaylo et al. 2021). To allow for a fungus to discriminate between self and nonself, fungi possess a robust multigenic system of heterokaryon incompatibility, based on *het* genes (genes which prevent formation of heterokaryons), which trigger cell death when gene products from two alleles are found in the same cytoplasm (Glass and Dementhon 2006). Within a fungal population these *het* genes are highly polymorphic and segregation of the various alleles results in each fungal individual being virtually guaranteed to have a distinct mix of *het* gene alleles compared to a different strain (Nauta and Hoekstra 1994; Czárán et al. 2014; Gonçalves and Glass 2020).

A common observation of such *het* genes is that they appear to be under balancing selection. First shown in *het*-c of *Neurospora crassa*, this balancing selection results in alleles being found in even frequencies in the population of this fungus (i.e. two alleles found at a 1:1 ratio, or three alleles at 1:1:1) (Wu et al. 1998). This pattern of even allele frequencies appears to be quite general, and has subsequently been found in other fungal species (Bastiaans et al. 2014; Milgroom et al. 2018; Ament-Velásquez et al. 2022). While it can be difficult to disentangle the cause of balancing selection (Spurgin and Richardson 2010), the generally accepted cause for nonself recognition genes such as *het* genes is negative frequency dependent selection (Nauta and Hoekstra 1994; Muirhead et al. 2002). This relationship between fitness and allele frequency emerges as a result of *het-* gene action being dependent on the alternate allele. As an allele becomes common in a population, more individuals will share this allele and this allele will be less useful to distinguish other individuals. Conversely, individuals with the alternate allele, which is now rare, will be able to detect other individuals as they will carry the alternate allele. This prevents either allele from reaching fixation, and thus both alleles are maintained longer than would be expected by random processes, with alleles often shared between related species in trans-species polymorphisms (Charlesworth 2006). Such patterns have been found in the genera *Neurospora, Cryphonectria*, and *Podospora* (Wu et al. 1998; Milgroom et al. 2018; Ament-Velásquez 2020).

Through various cell mechanisms, fusion between two fungal individuals, generally differing at multiple *het* loci, leads to the death of the nascent fused cell (Garnjobst and Wilson 1956; Glass and Kaneko 2003). Some known *het* genes trigger death due to protein-protein interaction encoded by different alleles of the same genes (allelic), while other *het* gene reactions are based on protein products of alleles of separate tightly-linked genes (non-allelic) within a larger haplotype (Kaneko et al. 2006). The biochemical pathway leading to death of this cell fusion differs depending on the *het* gene involved. Some *het* genes utilize systems based on Nod Like Receptors (NLR-like), with strong parallels to the immune system of animals and plants (Uehling et al. 2017; Heller et al. 2018). The action of *het* genes can also be based on prion formation of certain proteins, converting the alternate protein product into a cytotoxic protein that destabilizes the plasma membrane (Debets et al. 2012; Saupe and Daskalov 2012). For many *het* genes, the mode of action remains unknown. In the *Sordariomycete* genera *Neurospora* and *Podospora* many *het* genes contain a protein domain of unknown function, termed the HET domain (Saupe and Glass 1997; Zhao et al. 2015). While this domain is found in other fungi, the relationship between the HET domain and functional *het* genes in other fungal lineages remains uncertain (Fedorova et al. 2008; Dyrka et al. 2014; van der Nest et al. 2014).

The fungus *Aspergillus fumigatus* is globally dispersed and primarily decays dead plant material. Its spores are inhaled daily by birds and mammals, generally presenting no concern to the health of the organism. However, when the immune system is suppressed, such as during chemotherapy, immune suppressive therapy following organ transplantation or co-infection with influenza or SARS-cov2 in intensive care units this fungus can invasively grow into the lung tissue, leading to a condition called invasive aspergillosis (Schwartz et al. 2020; Arastehfar et al. 2021; Arné et al. 2021; Chong and Neu 2021). *A. fumigatus* can also chronically colonize human lungs in Cystic Fibrosis (CF) or Chronic Obstructive Pulmonary Disease (COPD) patients, and the same clonal isolate can be recovered from a patient sequentially for years (Bhargava et al. 1989; de Valk et al. 2009). First-choice treatment for severe *Aspergillus* disease is the azole antifungal class targeting the fungal ergosterol pathways. Increasingly, antifungal resistance is found due to environmental exposure of the isolates prior to infection of a human host (Verweij et al. 2009). As a potential alternative to traditional drug targets, the manipulation of cell death processes has been proposed to manage fungal infections (Weaver 2013; Hardwick 2018; Kulkarni et al. 2019). The first step for such research would be the identification of the genes that trigger heterokaryon incompatibility, the *het* genes, in *A. fumigatus*.

The genetics of *het* genes in other *Aspergilli* has been investigated previously, although not at a molecular level. Experimental studies in *A. nidulans* and *A. heterothallicus* both showed that incompatibility segregated in the progeny of sexual crosses, and was due to the action of several loci, although this was before the genomic era and specific genes were not identified (Kwon and Raper 1967; Anwar et al. 1993; Dales et al. 1993; Coenen et al. 1994). Using bioinformatic tools, sequence divergence combined with gene function information has been used as criteria to identify putative *het* loci in *A. fumigatus* (Fedorova et al. 2008). These diverged regions were termed “islands” of diversity in the *A. fumigatus* genome, with a highly divergent region flanked by large stretches of almost identical sequence. More recently, a reverse genetics study of genes encoding a HET domain in *A. oryzae* showed that expression of alternate alleles of most genes with a HET domain did not lead to an incompatible reaction, although some led to a modest reduction in growth (Mori et al. 2019). Within *A. fumigatus* it has been shown that environmental strains are not capable of forming stable heterokaryons, indicating abundant heterokaryon incompatibility genes (Weaver 2013; Zhang et al. 2019). Currently, validated *het* genes remain unidentified both in *A. fumigatus* as well as the genus *Aspergillus* in general.

Here, we combine whole-genome sequence data of sexual progeny from two heterokaryon incompatible environmental strains with phenotypic interactions to map the (in)ability to complement two auxotrophic mutations and form stable heterokaryons. Our presumption is that the inability to complement is related to *het* gene allelic differences. We then investigate the putative function and evolutionary history of these loci across related species and finally, we consider how the exceptional recombination rate of *A. fumigatus* may affect the process of balancing selection.

## Results

### Heterokaryon compatibility testing between offspring and parental strains

To map the genes causing incompatibility between our parental strains, we used heterokaryon complementation of auxotrophic nitrate pathway mutations. In this method, two haploid strains with different nitrate-deficiency mutations (e.g. one *nir* and the other *cnx*) will show vigorous heterokaryotic growth on nitrate as a sole N-source when compatible, but not when incompatible, due to sustained cell fusion. We produced from each of the two parents (AfIR974 and AfIR964) a *nir, cnx* and *nia* mutant, from which we crossed the AfIR974 *cnx* and the AfIR964 *nir* mutants. Among the 193 offspring, the distribution of genotypes (42 *nir/*wt; 51 *cnx/*wt; 55 *nir*/*cnx;* 45 wt/wt) was not significantly different from the expected 1:1:1:1 Mendelian ratio for two segregating loci (χ^2^ test, p=0.56)

In total 148 offspring were of a genotypic class that allowed heterokaryon compatibility testing to *nia* mutant versions of the parents (42 *nir/*wt, 51 *cnx/*wt *and* 55 *nir/cnx*). From these, we recovered five successful combinations forming a heterokaryon (3.4%) after three days of incubation. Three offspring (#118, 189, 108) could form heterokaryons with the parent AfIR964 *nia*, and two offspring (#166, 142) were compatible with the parental AfIR974 *nia*. Offspring that were compatible with a parent were only compatible with one parent. To increase the number of compatible pairs recovered, we tested each of the 42 *nir* strains against each of the 51 *cnx* strains of the offspring, resulting in testing of 2142 sibling pairings. Of these, 166 pairs (7.2%) formed heterokaryons. Of these 166 pairs, visible heterokaryons developed within three days for 133 pairs, whereas for 33 pairs, heterokaryons were visible only after five days.

### Genetic Mapping

From the 166 heterokaryon compatible sibling pairings and the 5 compatible parental-offspring pairs, we could form 15 vegetative compatibility groups (VCGs) (Supplemental File 2). Using the assumption that VCGs will have identical alleles at all *het* genes, we compared the genome sequence of the strains in each VCG at 14,153 variant sites previously identified (Auxier et al. 2022). As genotypes are expected to be similar within, but not between, groups we used the Shannon’s Entropy metric to determine similarity (see Methods). Mapping using the 133 isolates forming heterokaryons after three days resulted in peaks on chromosomes 2, 5, 6 and 8, which we termed *hetA, hetB, hetC and hetD*, respectively (Figure 1A). Each of these peaks rose above the significance threshold of 0.567, the highest 5% of the permutations. For each of these peaks, the window of significant markers spanned approximately 40kb with up to 20 predicted protein coding genes (Figure 1B-E). Within the larger VCGs, we observed a polymorphism where some pairings formed heterokaryons within three days, while other pairings formed heterokaryons after only five days (Supp.offspring.genotypes.xlsx; Pairwise Interactions tab). To investigate the genetic basis of this, we split the VCGs into sub-VCGs and performed a similar Shannon Entropy analysis (Supplemental Figure 1). This analysis recovered a single region on Chromosome 6, which we termed *hetE*.

**Figure 1:**
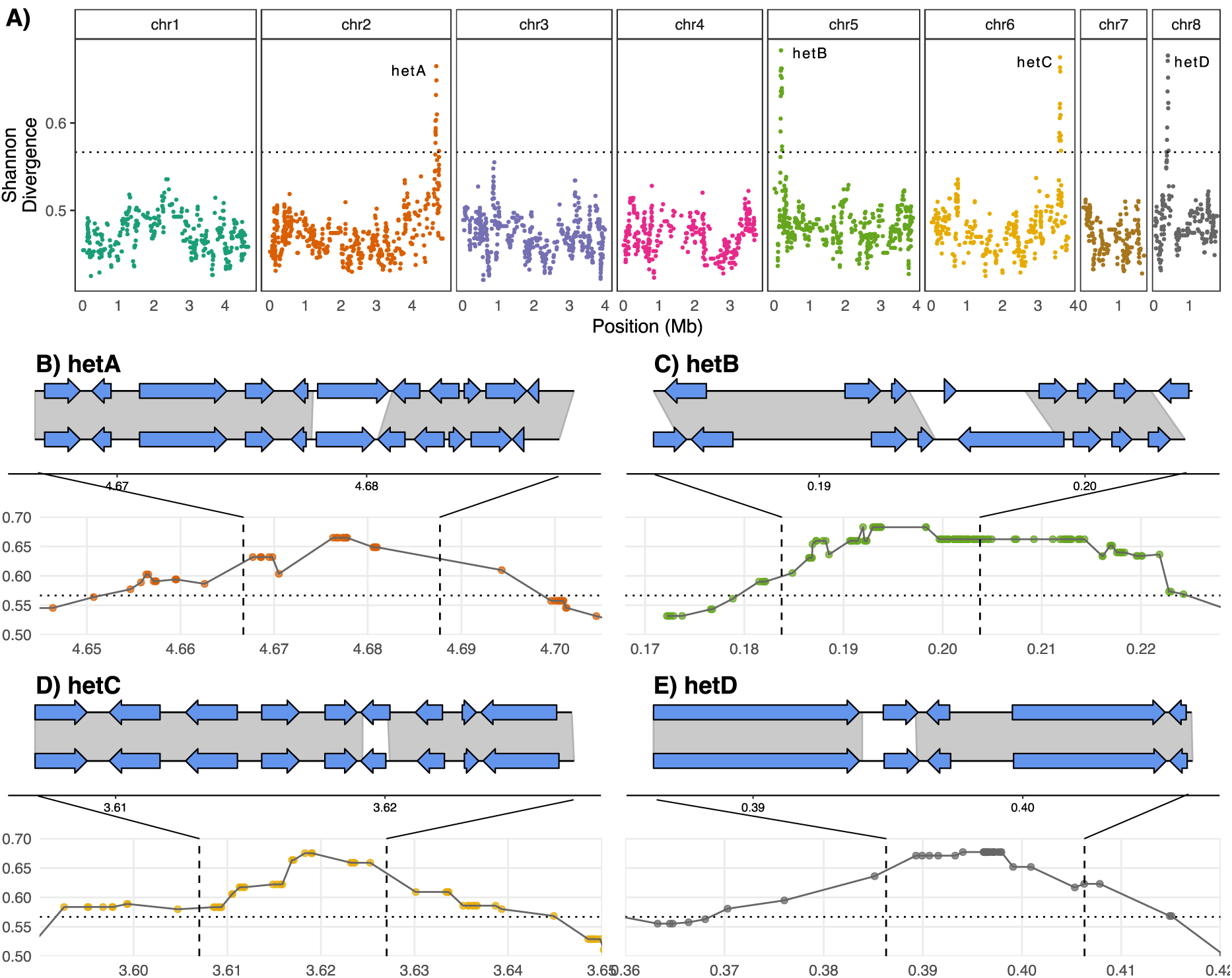
Mapping of *A. fumigatus het* loci based on heterokaryon-forming pairs. A) Genome wide mapping of heterokaryon compatibility after three days of incubation. Dots indicate variant positions, and associated Shannon Entropy phenotypic association. Significance threshold is determined by the top scoring 5% of 1000 random phenotypic permutations. B-E) Fine mapping of *hetA/B/C/D* loci. The lower part of each figure shows a closeup of the locus in A) that rises above the significance threshold. Dashed vertical lines indicate the variable genomic region with gene models shown with AfIR964 above AfIR974. Arrows indicate predicted gene models, and grey shading indicate regions with high sequence similarity between the two parental genomes. In these four loci, there is a region in the middle of the locus, with one associated gene, that is highly differentiated between parents, and lacks grey shading.

For each of the four *hetA/B/C/D* loci, the sequence similarity was high across the locus between the parents except near the centre of the locus with a single coding region with low sequence similarity (grey shading Figure 1B-E). For *hetA/C/D* this central divergent gene had two similarly sized alleles. For *hetB* we instead found that the two alleles had greatly differing coding sequence lengths, and in opposite directions. The *hetE* region was larger than the others, spanning approximately 80kb, with ∼40 predicted protein coding genes. The *hetE* locus contained three coding sequences in the AfIR974 allele, and four coding sequences in the AfIR964 allele, all of which had low sequence similarity between the parents (Supplemental Figure 1).

To verify that the five loci identified represented all the *het* loci segregating between these two parents, we backcrossed offspring predicted to differ at only one identified *het* locus from a parent. We then scored 40 offspring from each of nine such crosses, two independent crosses for *hetA/B/C/D* and one for *hetE*, for compatibility to the parental strain. In all crosses compatibility to the parent segregated as expected and did not significantly deviate from a 1:1 ratio (Supplemental Table 1).

### *het* Gene Identification and Validation

The predicted protein products of the *hetA* alleles both contained a nucleotide phosphorylase PNP_UDP, as well as a predicted nucleotide binding NB-ARC domain for the AfIR964 allele (Figure 2A). The *hetB* gene has two idiomorphic alleles, with the AfIR964 allele predicted to encode a 1327 amino acid protein that was predicted to contain a CHAT (Caspase HetF Associated with Tprs) domain, while the alternate *hetB* allele from AfIR974 was 149 amino acids long and had no predicted domains (Figure 2B). The alleles of the *hetC* gene were both predicted to produce a patatin-like protein of similar length (Figure 2C). The *hetD* alleles were also both predicted to encode similar sized proteins, both with a single PNP_UDP phosphorylase domain (Figure 2D). In contrast to the other *het* genes, the *hetE* locus contained multiple candidate genes all of which were polymorphic between parents (Figure 2E). Of these, one gene lacked identifiable domains, while another was the ortholog of a *A. nidulans* characterized gene *rosA* (See Discussion) and one was an uncharacterized C6 transcription factor (Figure 2E). The final candidate gene in the *hetE* locus contained a NACHT domain as well as Ankyrin-repeats and was found in the AfIR964 allele and not present in the AfIR974 allele. Comparison with Af293 showed an alternate allele of the NACHT/Ankyrin gene with low sequence similarity (Figure 2E).

**Figure 2:**
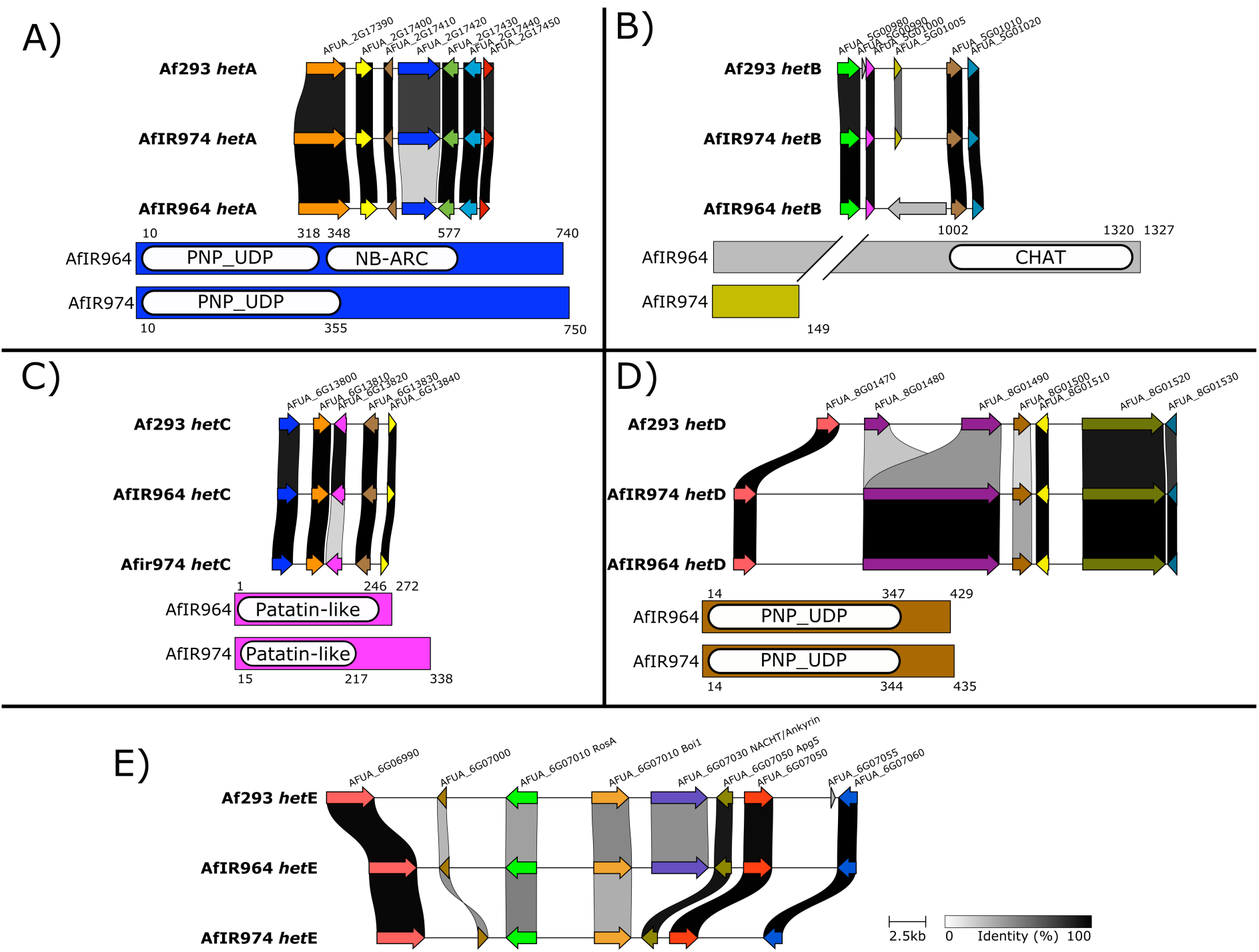
Genomic details of *het* genes. A-E) Top shows syntenic region of *het* gene with adjacent genes for AfIR974, AfIR964 and the reference strain Af293. Colours indicate gene identity, and links between genes indicate amino acid sequence similarity (legend in E). Below shows the protein sequence for each allele, with domains as predicted by InterPro. Colours of proteins reflect synteny analysis above.

We confirmed that the candidate *het* genes were causal to the phenotype using heterologous expression. Using a nuclear-localized AMA1 plasmid we cloned each allele including ∼500bp of flanking sequence for *hetA, hetB* and *hetC* (Figure 3A). We did not attempt validation of either *hetD* or any of the genes in the *hetE* locus. For each of the three loci tested, expression of the resident allele on the replicating plasmid led to no change in phenotype. However, introduction of the alternate allele led to an aberrant phenotype of abundant white mycelium and a complete absence of sporulation (Figure 3B-D). For both *hetA* and *hetC* both alleles had equal effect, with the same white mycelial phenotype. For *hetB*, expression of the *hetB2* allele in AfIR974 produced this same white non-sporulating phenotype, but the expression of *hetB1* in AfIR964 resulted in initial non-sporulating colonies that were not viable after transfer on selective media. Transfer of colonies to media without selection pressure allowed rapid sectoring, restoring the wild-type green sporulation, further indicating the phenotype resulted from the presence of the plasmid with the antagonistic *het* allele and not the transformation procedure (data not shown).

**Figure 3:**
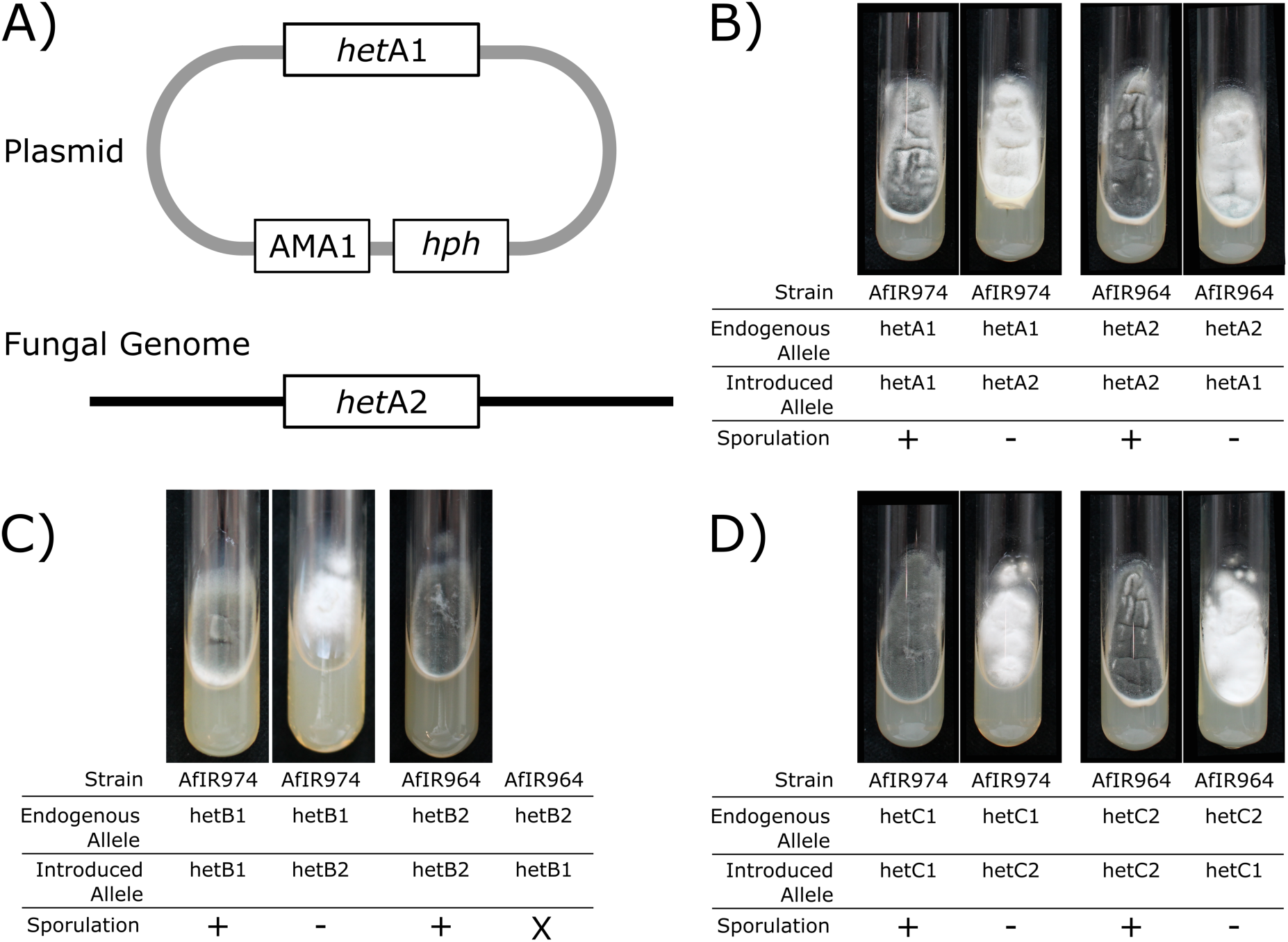
Validation of predicted *A. fumigatus het* genes using heterologous expression. A) Schematic of introducing an alternate *het* allele by transformation with the non-integrative AMA1 plasmid. B) Phenotypes of *hetA* transformants growing on hygromycin selective media, with transformants with endogenous alleles showing wildtype green sporulation, and transformants with the alternate allele with visible white mycelia due to lack of sporulation. C) Phenotypes as in B, but with transformants for *hetB* alleles. Note that AfIR964 with *hetB1* was not viable. D) Similar to B, but with transformants for *hetC* alleles.

### Balancing Selection

As heterokaryon incompatibility genes are expected to be under balancing selection, we assessed the strength of this selection across *A. fumigatus* in two datasets. Using genome-wide variants of 213 UK samples, as well as 178 German samples, we calculated Tajima’s D values across populations in sliding windows of 10 kb (Supplemental Figure 2). These two independent datasets had similar average Tajima’s D values (UK; average D = 0.05, Germany; average D = -0.31). Using window sizes of 10kb, we did not find increased Tajima’s D associated with any of our identified *het* genes nor with the mating-type locus (Supplemental Figure 2 A + B). *hetB* was the closest gene to a high Tajima’s D value, on chromosome 5, although this high D value is near the start of the chromosome, and careful inspection of the window surrounding *hetB* shows a D value below 2 for both population datasets.

As an alternate measure of balancing selection on these *het* genes, we reconstructed the phylogenetic relationships between amino acid sequences of *A. fumigatus* as well as the closely related *A. fischeri, A. lentulus* and *A. udagawae* for which multiple genome assemblies were available (Figure 4). For both *hetA* and *hetC*, the topology showed a trans-species polymorphism of two alleles, with each allele being found in all four species. The phylogeny of *hetD* showed a similar pattern of trans-species polymorphism, but instead with three alleles (Figure 4D). Of these three alleles for *hetD*, the D2 allele was present in all four species, but *hetD1* was not recovered in *A. udagawae*, while *hetD3* was not recovered from *A. fischeri*.

**Figure 4:**
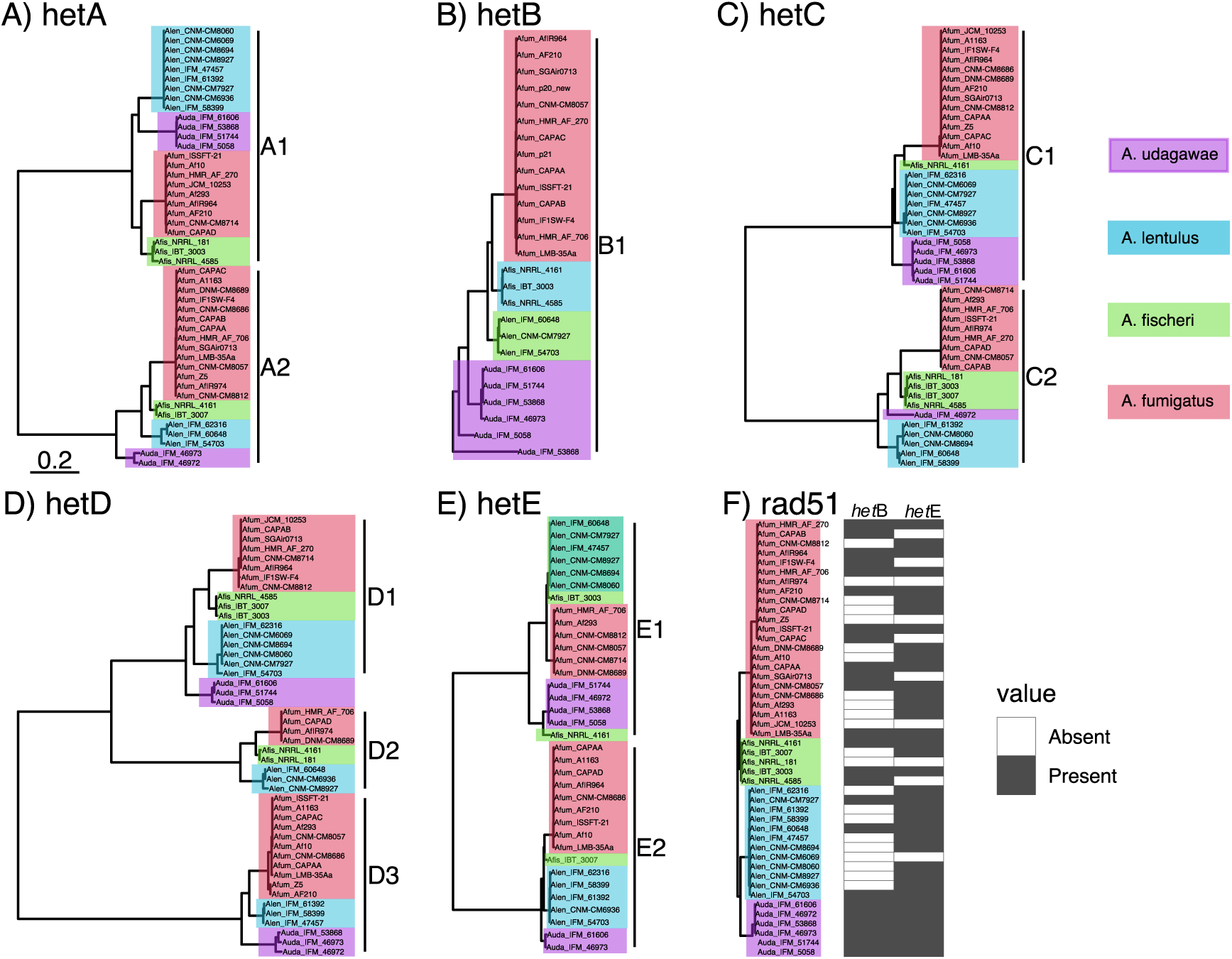
Phylogeny of predicted *het* protein sequences shows trans-species polymorphisms. Maximum likelihood unrooted phylogeny of amino acid sequence of related *Aspergilli*. Branch lengths indicate amino acid change, with tips coloured according to species. Salmon indicates *A. fumigatus*, green *A. fischeri*, light blue *A. lentulus*, and purple *A. udagawae*. Note that B) and E) only include the alleles with a protein present, and the absence alleles are indicated in F).

For *hetB* and *hetE*, which showed a presence/absence polymorphism between our parents (Figure 3), we reconstructed the phylogenetic relationships of the *het* alleles with the predicted coding sequence (Figure 4B and 4E) as well as recorded the status as presence/absence (Figure 4F). The *hetB* protease allele was present in approximately 50% (22/46) of the genome assemblies, and notably in all six *A. udagawae* assemblies. For *hetE*, the NACHT+Ankyrin repeat protein coding sequence was found in approximately 75% of genome (36/46), and the phylogeny showed two distinct protein coding alleles, meaning three alleles including the null allele, with a trans-species polymorphism. The absence allele was found in *A. fumigatus, fischeri* and *lentulus*. As a control analysis, alleles of the DNA repair gene *rad51* were separated between the species, with the alleles from each species being monophyletic (Figure 4F).

### Fine-scale population genetics of balancing selection in *A. fumigatus*

To attempt to reconcile the lack of population-level balancing selection with evidence for trans-species polymorphisms, we made use of public *de novo* genome assemblies from 300 previously sequenced individuals of *A. fumigatus*. Extracting the coding sequence and 5000 bp up- and downstream sequence allowed us to compare divergence between alleles of *hetA/B/C/D* as well as the *mat* locus. The Tajima’s D value was highest across the coding sequence, reaching values of >6 (Figure 5, top row). However, the drop-off to a neutral D value was abrupt, with values dropping below 2 within 500bp of the start/stop codon. This abrupt and rapid change was mostly mirrored in the nucleotide diversity within and between the allele classes (Figure 5, bottom row). An exception was found in the *mat* locus, where the nucleotide diversity was somewhat lower across the left-hand side of the coding region. Close inspection of the alignments revealed that this was due to a sharing of the terminal ∼300bp of the α-box protein.

**Figure 5:**
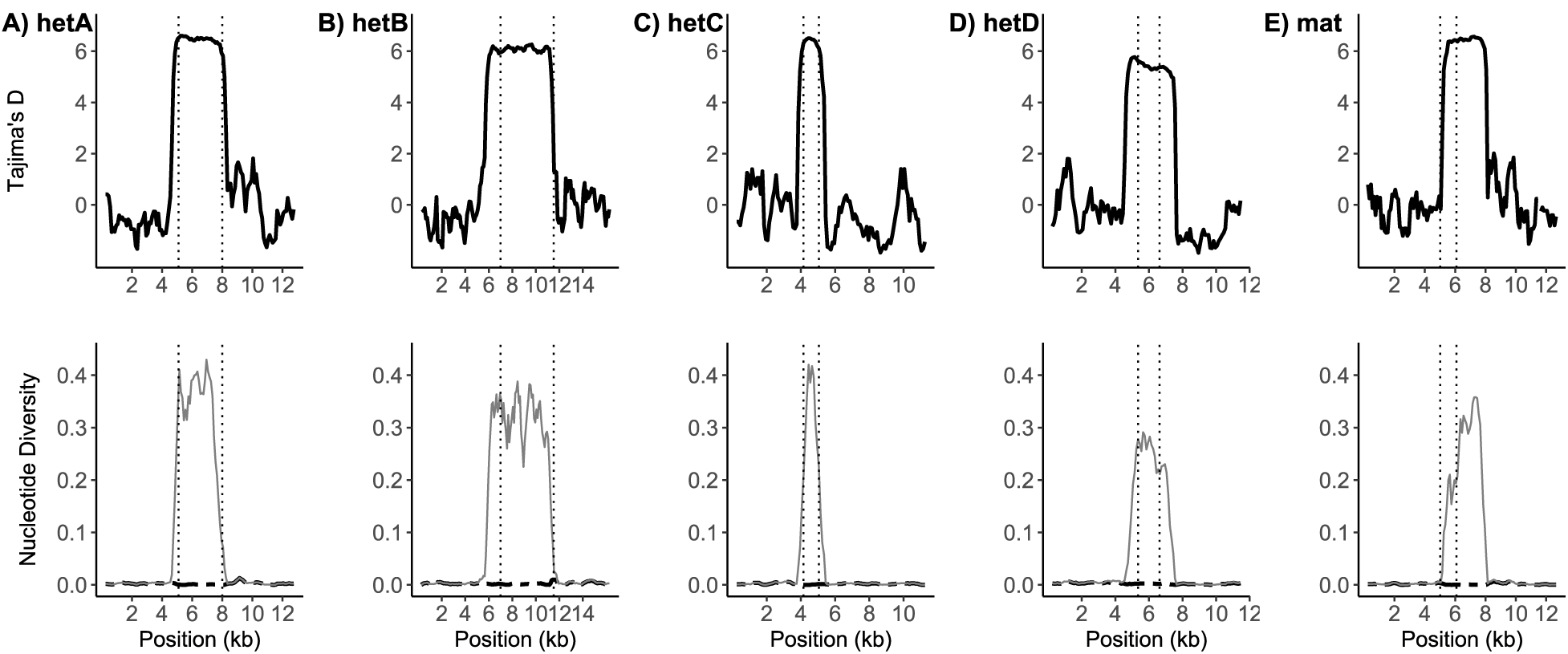
Signatures of balancing selection in *A. fumigatus* are highly restricted. Above is plotted the Tajima’s D value for the region of each of the *hetA/B/C/D* as well as the *mat* locus and 5kb of flanking sequence. Dashed vertical lines indicate the start and stop codon of the respective gene, based on the Af293 annotation. The nucleotide diversity is plotted below, with thin grey lines indicating the difference between alleles, and the thick black dashed line indicated nucleotide diversity within an allele.

## Discussion

It is assumed that fusion between conspecific individuals is a risky behaviour, and so it is rigorously policed in fungi. As such, heterokaryon incompatibility is expected to be common between strains of *A. fumigatus*, and has been reported previously (Weaver 2013; Zhang et al. 2019). Our results here reveal the polygenic nature of this heterokaryon incompatibility. For our two parents, both isolated in 2005 in Dublin, Ireland, five loci contribute to incompatibility. Using a single cross and approximately 150 offspring, we were able to map all responsible loci with high resolution, highlighting the power of genetic mapping in *A. fumigatus* with its high recombination rate (Auxier et al. 2022). These identified *het* genes are to our knowledge the first identified in any *Aspergillus* species, and the first for any member of the class *Eurotiomycetes*.

Two of these *het* genes, *hetC* and *hetD* are consistent with effector domains commonly found in NLR genes, but without the other components (Uehling et al. 2017). The alleles of *hetD* encode similarly sized PNP-UDP domains, while the alleles of *hetC* both encode a patatin-like domain. These patatin domains are also found in the patatin-like protein (PLP-1), which interacts with the SNARE domains of the nearby *sec*-9 gene in *Neurospora crassa* (Heller et al. 2018). However, the action of the gene products of *hetC* and *hetD* seems to occur in isolation, since there are no neighbouring genes with divergent alleles. It is interesting to note that in *N. crassa* there is another *plp* gene, *plp*-2, which also shows evidence of balancing selection, but non-allelic differences between *plp*-2 and *sec*-9 did not trigger cell death (Heller et al. 2018). As differences at a patatin-like protein appear sufficient to trigger heterokaryon incompatibility in *A. fumigatus*, perhaps there are also allelic interactions in *N. crassa plp*-2 that do not require a partner gene to trigger nonself recognition.

Alleles from two of the *het* genes we identify, *hetA and hetE*, have a more traditional NLR-like structure, common to fungal *het* genes as well as immune-system genes in plants and animals (Uehling et al. 2017). Differences at *hetA* completely block heterokaryon formation, while heteroallelism at *hetE* delays, but does not prevent, heterokaryon formation. This is strikingly similar to the “partial” *het* genes previously identified in *A. nidulans* (Coenen et al. 1994). There multiple alleles were identified that caused delayed heterokaryon compatibility, however differences at multiple of these “partial” *het* genes was sufficient to fully block heterokaryon formation (Coenen et al. 1994). It would be interesting to test the interaction between alternate alleles of *hetE* both encoding NLR proteins, and whether this interaction would also only delay or completely block heterokaryon formation. Across *A. fumigatus* and related species, we recovered two different NLR-encoding alleles for *hetE*, and one allele without a coding sequence. This situation, with two alternate protein coding alleles and one null allele, is similar to the ABO blood group system in primates where even the null allele shows trans-species polymorphism (Ségurel et al. 2012). However, it is possible that the likely candidate gene, producing the NACHT+Ankyrin repeat protein, is not acting alone. Adjacent to this gene is AFUA_6G07020, an ortholog of *boi1*. This gene is known from yeast to function in vesicle fusion during exocytosis (Kustermann et al. 2017: 1; Masgrau et al. 2017: 1). Interestingly, a similar pairing has recently been described in the *het* genes of the plant pathogen *Botrytis cinerea*: *Bcvic1*, a NACHT+Ankyrin encoding gene, and *Bcvic2*, which encodes a SNARE syntaxin protein (Arshed et al. 2022). These SNARE domains are also known from the *sec*-9/SNARE system in *N. crassa*, and *hetQ* of *Podospora anserina*, which both involve an NLR type protein and a SNARE protein (Heller et al. 2018). It may be that *hetE* of *A. fumigatus* has an interaction with neighbouring *boi1* in an analogous role. Notably, the *sec*-9/*plp*-1 interaction in *N. crassa* has a strong phenotype at the germling stage, but not in developed mycelia (Heller et al. 2018). This may point to a stronger phenotype for *het*E here, if tested during germling fusion. Recently, methods for visualizing germling fusion have been developed which may assist in further studies (Macdonald et al. 2019).

The *hetB* locus in *A. fumigatus* presents two alleles that share no protein similarity, idiomorphs, with one allele encoding a large protease, and the other allele a short protein without annotated domains. This large protease is predicted to contain a CHAT domain, known to function in mammalian apoptosis, and found in other metacaspases in non-mammalian lineages (Hoffman et al. 1997; Aravind and Koonin 2002; Bouchier-Hayes and Martin 2002). This domain has not previously been identified to be involved in heterokaryon incompatibility in a fungal lineage. The involvement of a CHAT domain raises interesting questions about downstream pathways since fungal cell death is not considered homologous to mammalian apoptosis (Hardwick 2018; Kulkarni et al. 2019). The mechanism behind this protease triggering incompatibility may be related to a recent report of proteolytic cleavage leading to cell death described in the *het-*Q system in *Podospora anserina* (Clavé et al. 2022). Verification of the biochemical target of the protease of *hetB*, whether it is produced by the small coding sequence of the alternate allele, is an important next step.

A conspicuous absence in our data is an association of the mating-type locus and heterokaryon incompatibility. In an obligately outcrossing species, the mating-type locus experiences balancing selection due to frequency-dependent selection. As an increased number of loci under balancing selection across the genome comes with a cost of reduced genetic diversity at other locations (Wittmann et al. 2022), it may seem logical that such a locus would also be involved in heterokaryon incompatibility, which has been previously suggested (Crozier 1986). This situation is found in many fungal species such as *N. crassa*, where the mating locus also produce heterokaryon incompatibility (Newmeyer et al. 1973). Even within *Aspergilli* the mating locus in outcrossing *A. heterothallicus* was genetically very tightly linked with *het* gene activity, although the molecular genetics are unknown (Kwon and Raper 1967). However, the absence of mating-type associated heterokaryon incompatibility has also been observed, for example in *Cryphonectria parasitica* (McGuire et al. 2004). A by-product of this absence of mating-type associated incompatibility in *A. fumigatus* may be the formation of a sexually fertile diploid heterozygous for the mating type. This has been recently suggested in the diploid formation model of *A. latus*, although this model did not incorporate the difficulties imposed by heterokaryon incompatibility between wild isolates (Steenwyk et al. 2020). While diploids have been previously observed in clinical isolates of *A. fumigatus*, these diploids were almost completely homozygous having arisen by genome duplication, avoiding incompatibility (Engel et al. 2020). Regarding the mating locus itself, population level analysis showed that the effects of balancing selection was not as sharp as expected. This is due to a partial sequence of the transcription factor of the MAT-2 allele being also found in the MAT-1 allele, as reported previously (Paoletti et al. 2005).

Of the loci described here, only for *hetE* are non-allelic interactions, the interactions between alleles of linked genes instead of homologs, probable. In other species studied in detail, such non-allelic interactions seem common: *het-*R/V and *het*-C/D/E in *Podospora* (Saupe et al. 1995; Saupe 2000), the *hetC*/*pinC* and *het*-6 systems in *Neurospora* (Smith et al. 2000; Kaneko et al. 2006), the *Bcvic1/Bcvic2* system in *Botrytis*, and both the *vic2* and *vic6* loci in *Cryphonectria* (Choi et al. 2012). The extremely high recombination rate in *A. fumigatus* (1% of offspring will be non-parental across a distance of 2.5kb) (Auxier et al. 2022) may mean that such non-allelic interaction present a fitness cost. Any offspring where there was a crossover between the two genes in a non-allelic interaction would then contain an incompatible pair of alleles, and would trigger non-self-recognition within its own cytoplasm (Chevanne et al. 2009). However even for *hetE*, which may have non-allelic interactions with *boi1*, there is a large region of high sequence divergence, which is likely restricting recombination, as the meiotic machinery requires stretches of identical sequences during the initial homology search (Li et al. 2006). Therefore, the fact that *hetE* sits within a large divergent locus may itself prevent crossovers that would otherwise lead to self-incompatible offspring.

A surprising finding of this study was the lack of population level signatures of balancing selection when calculated using window sizes common to such studies (Heller et al. 2018; Ament-Velásquez 2020). It is well established that the mating locus as well as *het* genes are under negative frequency dependent selection, leading to balanced polymorphisms. Our finding of extensive trans-species polymorphism confirmed that balancing selection was acting as expected for these alleles. However, we failed to find a signature with standard population genomics analysis. It appears this discrepancy can be reconciled through a closer analysis of genotypes. Genome-wide studies typically use blocks of 10 kb or higher to average across the genomic stochasticity of individual variants. However, this assumes that the signal from balancing selection, like any other selection, will be carried partially to the neighbouring sequence, the “footprint” of selection, due to linkage with the selected locus. In *A. fumigatus*, we see that variation between allelic classes only extends ∼200bp up- or downstream of the coding regions. This means that standard windowed analysis for this species is likely to skip over strong signals of selection, whether from balancing selection or otherwise. Compounding this narrow footprint of balancing selection, the separation between these alleles has existed for so long that the sequences have diverged to a point that standard genomic methods fail. Within such regions, typical short-read DNA data will not be able to be mapped to the syntenic region between isolates. Downstream analysis will then generally remove such variant sites, due to “missing” data.

The *het* genes we identified appear to have been under balancing selection for several millions of years at minimum. We find shared alleles across a group of four related species. However, we were limited in our sampling to species with multiple available genome assemblies, and thus closely related species like *A. oerlinghausenensis* could not be used (Houbraken et al. 2016), and neither more distant species like *A. clavatus*. We would expect that the alleles for these five genes are shared across an even wider number of species, when additional genome assemblies become available. Across the species studied, we found that the allelic diversity was limited to a region only ∼250bp outside the coding region of the gene. This is likely related to the high recombination rate in this species, reducing linkage to nearby regions. This observation has been described before as “genomic islands of divergence”, but the cause was unclear at the time (Fedorova et al. 2008). Although there are few genome-wide empirical analyses of negative frequency dependent selection, it seems that generally the window of balancing selection is wider than observed here. A recent analysis of balancing selection on the S gametic self-incompatibility locus showed that increased nucleotide diversity was seen several tens of kilobases from the selected locus (Veve et al. 2022). Recent analysis of the fungus *Podospora anserina* showed that signatures of balancing selection could be determined from population level analysis looking at 10kb windows (Ament-Velásquez 2020). However, the general relationship between recombination rate and the size of the footprint is unclear, as the divergent region surrounding the *MLO2b* immunity gene in *Capsella sp*. is also tightly restricted, despite a much lower rate of recombination (Sicard et al. 2011; Koenig et al. 2019). The interaction between the age of balancing selection, recombination rate, and the resulting window of sequence divergence requires future study.

Fungal heterokaryon incompatibility involves a sort of paradox. It is understood to be ancestral at least to Dikaryotic fungi, and present in all extant filamentous species. Yet there are no ubiquitous fungal “*het*” genes. Thus, each species, or set of related species, require independent study to understand the repertoire of fungal *het* genes (Paoletti 2016). The five loci we identify here are unlikely to be the only variable *het* genes in the species, and analysis of additional isolates of this species likely will reveal additional *het* genes. However, detailed understanding of additional nonself recognition responses is still only an initial step in understanding heterokaryon incompatibility. The mechanism by which these allelic variants trigger cell death remains largely unknown and requires further study. Deciphering the programmed cell death pathway in this species may present novel targets for treating human aspergillus infections and may open the way to eliminate incompatibility barriers to parasexual recombination for improvement of industrially important asexual *Aspergilli*.

## Materials and Methods

### Media

For testing of auxotrophic mutants and heterokaryon compatibility testing, Minimal Medium (MM) was used. MM consists of 6.0 g NaNO_3_, 1.5 g KH_2_PO_4_, 0.5 g MgSO_4_. 7H_2_O, 0.5 g KCl, 10 mg of FeSO_4_, ZnSO_4_, MnCl_2_, and CuSO_4_ and 15 g agar per 1000 mL H_2_O (pH 5.8) (Zhang et al. 2019). Alternative nitrogen sources (nitrate, nitrite and hypoxanthine) were added at 5 mM as necessary. MM supplemented with urea (0.3g/L) was permissive for all mutant growth and was used for general culturing of auxotrophic mutants and sexual offspring.

### Recessive markers introduction in the parental isolates

To allow for heterokaryon testing, recessive auxotrophic markers (nitrate non-utilizing mutations *nia, nir* and *cnx*) were isolated from spontaneous mutations in spore suspensions of parental isolates AfIR974 and AfIR964. To isolate these spontaneous mutations, spore suspensions (approximately 1×10^6^ spores) were spread on MM medium supplemented with 200 mM of chlorate and 5 mM urea. As chlorate is toxic to wild-type *A. fumigatus* this method selects for strains deficient in nitrogen metabolism. After three days of growth at 37 °C chlorate resistant sectors were isolated, purified and classified as *nia, nir* or *cnx* mutants based on differential N-source media tests (Cove 1976).

### Recessive markers segregation of sexual offspring

The sexual cross was made from parental isolates AfIR964*cnx* and AfIR974*nir*, which were co-inoculated on Oatmeal Agar followed by incubation at 30 °C. The genotypes of the offspring were determined for a previous study (Auxier et al. 2022). Offspring were phenotyped for *cnx* and *nir* markers as above.

### Heterokaryon compatibility testing

Heterokaryon compatibility of offspring was tested by co-inoculating conidia (approximately 10^5^) of isolates with complementing *nir, nia* or *cnx* markers on MM with NO_3_ as the sole nitrogen source (Zhang et al. 2019). All possible combinations of single mutant offspring with complementing *nir* and *cnx* markers were tested for compatibility. In addition, all *nir, cnx* and *nir/cnx* offspring were tested for compatibility with *nia* mutants of parental isolates. Following co-inoculation, plates were incubated at 37 °C, and heterokaryon formation, vigorous growing yet irregularly shaped colonies, was recorded after three and five days of incubation. Pairings that successfully formed heterokaryons were re-tested in triplicate, with additional testing of spore mixtures on MM+NO_3_ to ensure heterokaryotic growth was not due to contamination.

### Segregating crosses

Segregation of individual *het* loci was tested by backcrossing offspring that differed from one of the parents at only one *het* locus, (i.e. shared alleles for *hetA,C,D,E* but not *hetB* for example), and with the opposite mating type. Crosses were performed on Oatmeal Agar as above, and sexual offspring were isolated from cleistothecia following a 1 hour 70 °C heatshock treatment to kill asexual spores. Only *cnx* offspring were used, 40 of which were tested for compatibility against the *nia* version of the parental isolate as above.

### *het* loci association mapping

To map the regions associated with incompatibility, the 166 offspring and 2 parents were split into 15 compatibility groups based on heterokaryon testing. For each of the 14,153 bi-allelic markers in this mapping population identified previously (Auxier et al. 2022), the genetic similarity within a group was tested using the combined Shannon Entropy of the allele frequency within a group. As each group should be defined by genetic similarity within a group for a set of loci, Shannon Entropy allows to test whether the genotypes in a group are informative for a phenotype, without influence of the variation in compatibility group size. As the markers were all biallelic, the formula was:

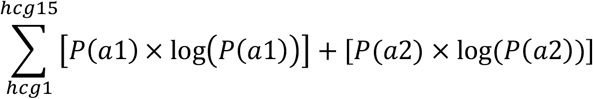

Where hcg1 to hcg15 are the different heterokaryon compatibility groups, and P(a1) and P(a2) are the allele frequencies of the two alleles within each group. Thus, the association for a marker is the summed Shannon entropy for the 15 recovered compatibility groups. To compute a null distribution, the genotypes of the compatible groupings of the 166 offspring were randomly distributed into 1000 replicate populations with compatibility group sizes the same as our actual results. Threshold cut-off was determined on the 95 % of the distribution. Fine-mapped regions were visualized with gggenes and clinkr (Gilchrist and Chooi 2021; Hackl and Ankenbrand 2022), and domains of candidate genes annotated with InterProScan (Paysan-Lafosse et al. 2022).

### *het* gene confirmation

To validate our predicted *het* genes as causal to the incompatible phenotype, we expressed the alternate alleles from autonomously replicating nuclear AMA1 plasmids. First, we PCR-amplified the predicted gene from gDNA of either parent using BioVeriFi polymerase (PCR Biosystems Inc, Pensylvannia USA; PB-10.42-05), including the upstream and downstream intergenic space (Supplemental Table 2). To include the native regulatory elements, we included approximately 750 bp of flanking regions, in some cases up to ∼50 bp of the adjacent gene. The plasmid backbone was also amplified with BioVeriFi polymerase and ligated with the insert using NEBuilder HiFi DNA Assembly Master Mix (New England Biolab Inc. #E2621L) according to manufacturer directions and transformed in to Mach1 competent *E. coli* cells (ThermoFisher Scientific Inc.). Plasmid was then extracted from these cells, and ∼1 μg was used in PEG-mediated protoplast transformation (van Rhijn et al. 2020). Transformed colonies were selected based on growth on Malt Extract Agar (20 g/L) + 300 μg/mL Hygromycin B (H0192; Duchefa Biochemie, Haarlem, the Netherlands).

### Cross-species comparison

To assess the extent of trans-species polymorphisms, we made use of publicly available genome assemblies (Supplemental File 1). We used tblastn v2.8.1 to search the genomes of related *Aspergillus* species using predicted protein alleles from AfIR964, AfIR974, and Af293 (Camacho et al. 2009), using an e-value cutoff of 1e-75. To avoid partial matches based on conserved domain, we filtered results to be >300bp long. The corresponding DNA sequences were extracted, and aligned with mafft v7.427 using the --auto command (Katoh and Standley 2013). Phylogenetic relationships were inferred using iqtree v1.6.1using default settings (Nguyen et al. 2015). The resulting most likely tree was midpoint rooted and visualized with ape and ggtree (Paradis et al. 2004; Yu et al. 2017).

### *A. fumigatus* population genome scan

To assess the effects of balancing selection surrounding genes expected to be under balancing selection, we used data from two resequencing studies (Barber et al. 2021; Rhodes et al. 2022). From the UK dataset of Rhodes et al., we downloaded the VCF file from the publication, while for the Barber et al. dataset we called variants as described previously (Auxier et al. 2022). Tajima’s D values were calculated in windows sizes of 10,000 bp using vcftools (Danecek et al. 2011).

### *het* gene diversity

To determine the nucleotide diversity across the *A. fumigatus* species, we made use of genome assemblies from a recent large population resequencing study (Barber et al. 2020; Barber et al. 2021). The genome assemblies of the *A. fumigatus* strains used for the trans-species polymorphism analysis (see above) were added to this dataset, making a total of 334 *A. fumigatus* genome assemblies. For each of *hetA/B/C/D*, as well as the mating type as a positive control, we extracted the corresponding gene from Af293, as well as 5 kb of flanking sequence. Due to the high sequence divergence of the *het* alleles, we used the high sequence similarity of flanking regions to isolate homologous sequences. This sequence was used with BLASTN to search the genomes of each assembly, and the region inside the borders of the flanking sequence was retained. The sequence was extracted using samtools, and aligned using mafft v7.427 using the GINSI algorithm as we expected low similarity in the centre of the alignment (Li et al. 2009; Katoh and Standley 2013).

## Acknowledgments

This work was supported by the Netherlands Research Organization (NWO ALGR.2017.010 to BA; ZonMV 451001022 to ES). We thank Drs. Sandra Lorena Ament-Velásquez and Ivain Martinossi-Allibert for valuable discussion regarding fungal balancing selection. We thank Wendy Adriaens for assistance with the *Aspergillus* trans-species polymorphisms. We particularly thank Ke Luan for assistance with phenotyping of the heterokaryon compatibility pairings.

## Data and material availability

Strains and plasmids are available by contacting corresponding authors. Code and data for analyses used in this manuscript is available at: https://github.com/BenAuxier/asp_fum_het

## Author Contributions

Conceptualization: AJMD, BA and ES. Experimental work: JZ, KL, KS, FRM, BA. Bioinformatic analysis: BA, JvdH. Manuscript draft: BA. Manuscript revisions: BA, AJMD, DA, ES, JZ. All authors read and approve publication of the manuscript.

## Supplementary Data

**Supplemental Figure 1:**
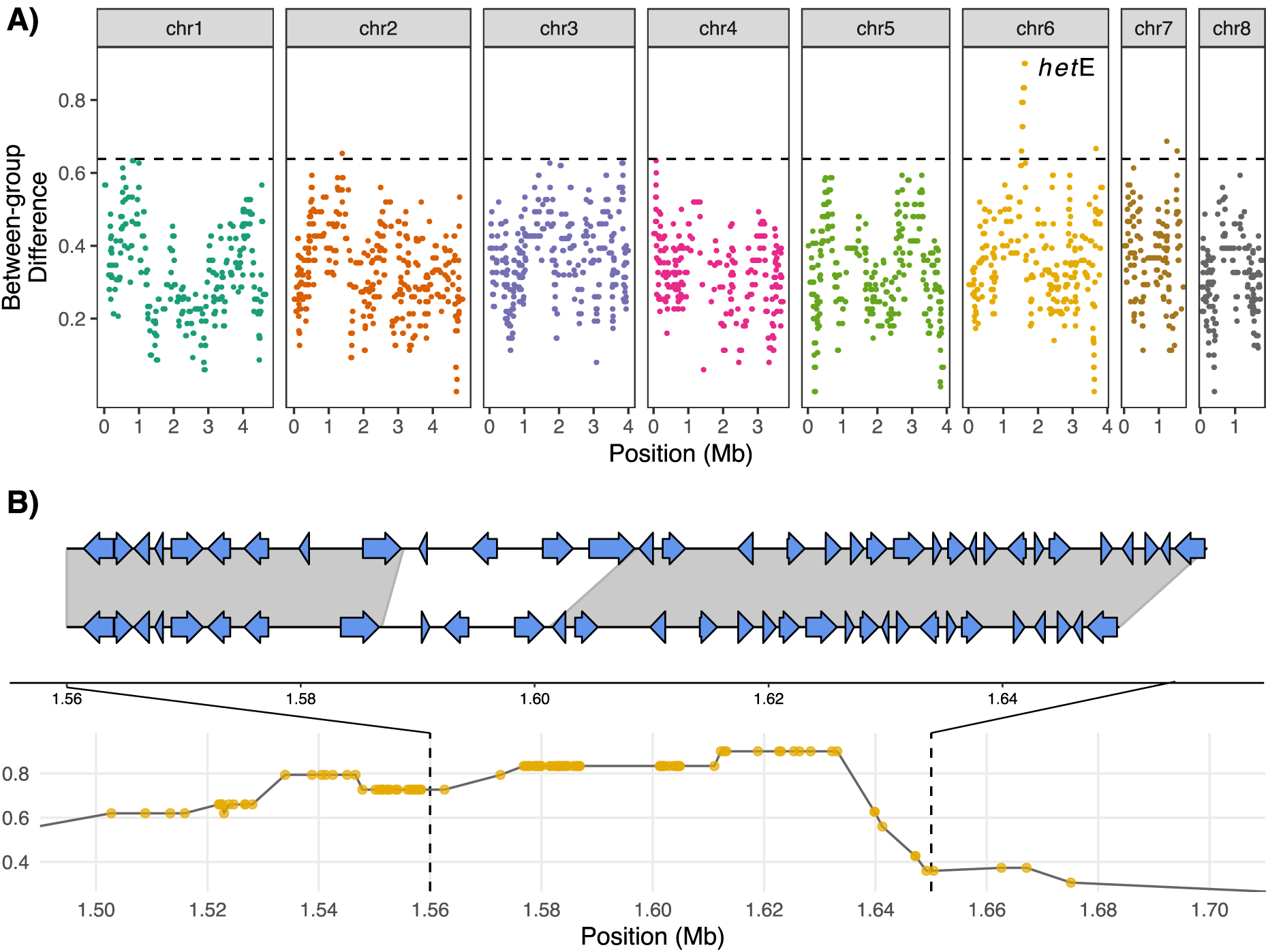
Genetic mapping of locus controlling compatibility after five days. Identification of *hetE* locus based on difference between heterokaryon formation at three vs. five days. A) Genome wide plot of allelic differences between 3-day group and 5-day group (See Methods). The peak on Chromosome 6 is indicated as *hetE*. B) close up of *hetE* peak similar to that found in Figure 2A. Note that the divergent window for *hetE* covers several predicted coding genes.

**Supplemental Figure 2:**
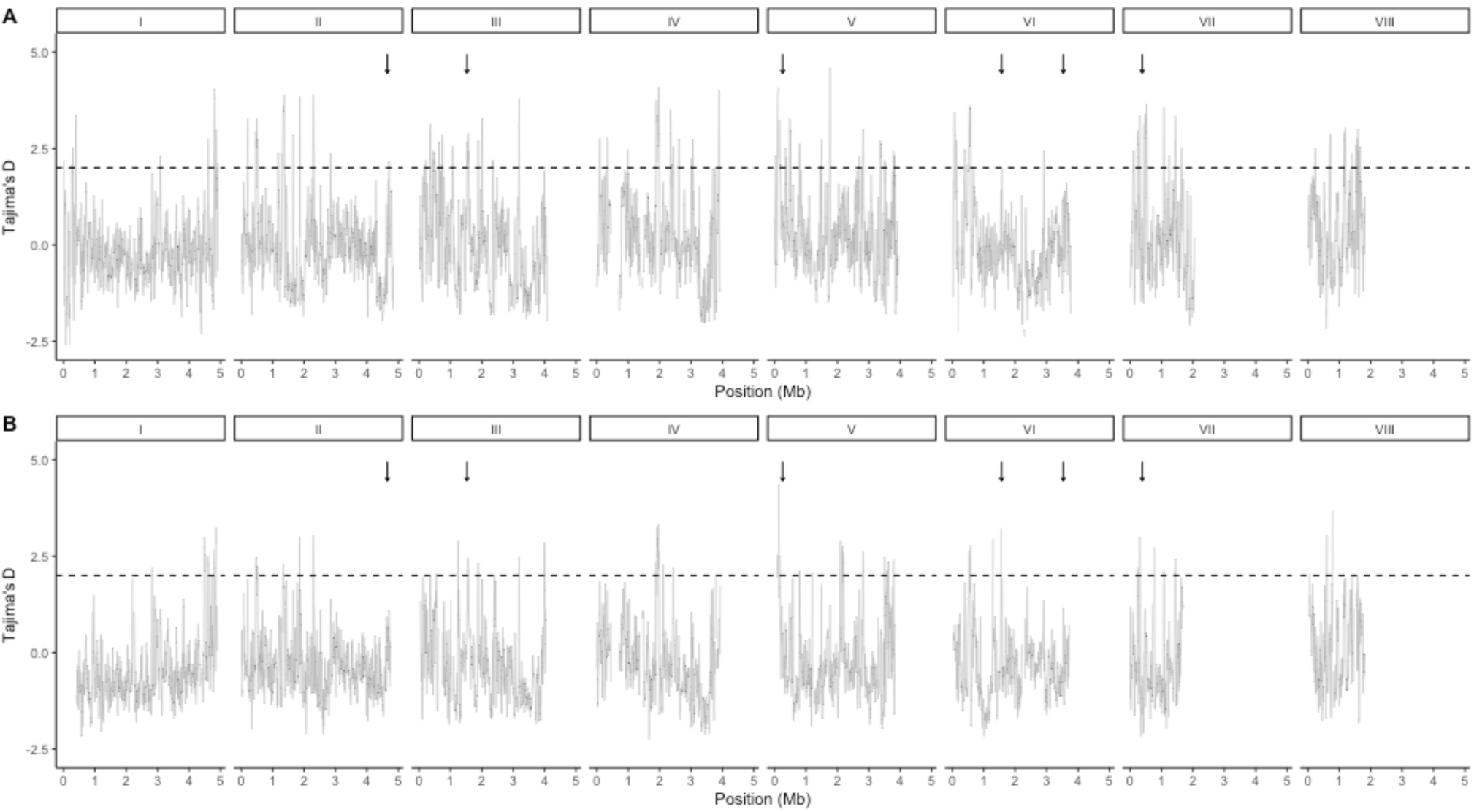
Genome-wide balancing selection in *A. fumigatus*. A) shows data from VCF file consisting of 213 UK samples (Rhodes et al. 2022). B) shows data computed from German samples described in (Barber et al. 2021) consisting of 178 German samples. For both populations, Tajima’s D is shown in 10kb windows, and an arbitrary threshold of 2 is applied as a dotted horizontal line. Arrows indicate position of *hetA, mat, hetB, hetC, hetE*, and *hetD*, in that order.

**Supplemental Table 1:** Data from 6 segregating crosses to test identified loci.

| parent1 | parent1<br>geno | mat | parent2 | parent2<br>geno | segregating | Spores<br>tested | Compatible with<br>parent | $\chi^2$ value | P<br>value |
| --- | --- | --- | --- | --- | --- | --- | --- | --- | --- |
| 192 | 11101 | MAT-1 | AflR964 | 11111 | hetD | 40 | 20 | 0 | 1 |
| 215 | 10111 | MAT-1 | AflR964 | 11111 | hetB | 34 | 18 | 0.11 | 0.73 |
| 71 | 11110 | MAT-1 | AflR964 | 11111 | hetE | 40 | 18 | 0.4 | 0.52 |
| 66 | 1000 | MAT-2 | AflR974 | 00000 | hetB | 40 | 19 | 0.1 | 0.75 |
| 171 | 11011 | MAT-1 | AflR964 | 11111 | hetC | 24 | 11 | 0.16 | 0.68 |
| 169 | 10000 | MAT-2 | AflR974 | 00000 | hetA | 40 | 22 | 0.4 | 0.5<br>2 |
| 154 | 11110 | MAT-1 | AflR964 | 11111 | hetE | no cross | no cross | N/A | N/A |
| 140 | 10111 | MAT-1 | AflR964 | 11111 | hetB | no cross | no cross | N/A | N/A |

**Supplemental Table 2:**
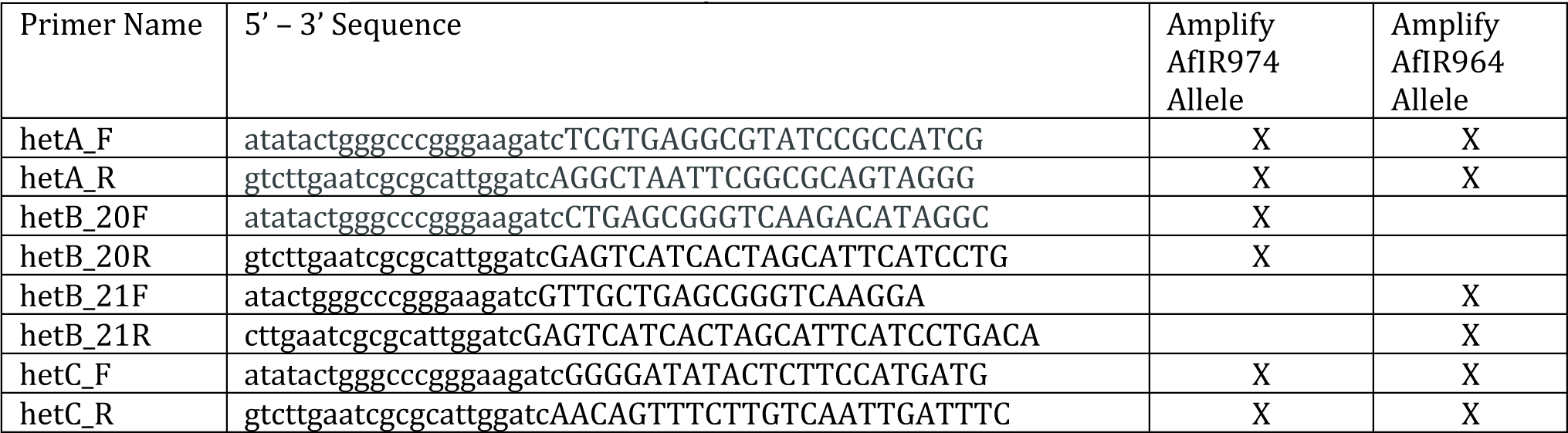
Primers used to construct *het* AMA1 plasmids

